# Bone Marrow Stromal Cells in a Mouse Model of Metal Implant Osseointegration

**DOI:** 10.1101/2020.08.17.254433

**Authors:** Alexander Vesprey, Eun Sung Suh, Didem Göz Aytürk, Xu Yang, Miracle Rogers, Branden Sosa, Yingzhen Niu, Ivo Kalajzic, Lionel B. Ivashkiv, Mathias P.G. Bostrom, Ugur M. Ayturk

## Abstract

Metal implants are commonly used in orthopaedic surgery. The mechanical stability and longevity of implants depend on adequate bone deposition along the implant surface. The cellular and molecular mechanisms underlying peri-implant bone formation (i.e. osseointegration) are incompletely understood. Herein, our goal was to determine the specific bone marrow stromal cell populations that contribute to bone formation around metal implants. To do this, we utilized a mouse tibial implant model that is clinically representative of human joint replacement procedures. Using a lineage-tracing approach with the *Acta2.creERT2* and *Tmem100.creERT2* transgenic alleles, we found that *Pdgfra*- and *Ly6a*/*Sca1*-expressing stromal cells (PαS cells) multiply and differentiate in the peri-implant environment to give rise to osteocytes in newly formed bone tissue. Single cell RNA-seq analysis indicated that PαS cells are quiescent in uninjured bone tissue; however, they express markers of proliferation and osteogenic differentiation shortly after implantation surgery. Our findings indicate that PαS cells are mobilized to repair bone tissue and facilitate implant osseointegration following surgery. Biologic therapies targeting PαS cells might improve osseointegration in patients undergoing orthopaedic procedures.

## INTRODUCTION

More than 6 million orthopaedic surgeries are performed in the United States every year. Approximately 1 million of these are knee or hip replacement procedures, and the frequency of these surgeries is expected to increase by more than 70% over the next decade [1]. Yet, up to 15% of patients undergoing joint replacement surgery develop complications and require complex and costly revision surgeries [2]. A leading cause of implant failure is aseptic loosening [2], which results from the inability of native bone tissue to sufficiently bond with the implant (i.e. osseointegration). An improved understanding of bone tissue’s response to injury that occurs during joint replacement surgery, and of mechanisms that promote implant osseointegration, could help devise novel biologic therapies that will reduce implant failure.

Multiple stromal cell populations exist in the long bone marrow of the appendicular skeleton that could give rise to osteoprogenitor cells that participate in osseointegration [3, 4]. One such population is identified by the expression of cell surface marker proteins PDGFRA and Sca1 (also referred to as PαS cells), and is predominantly located around arteriole-type blood vessels [5, 6]. PαS cells can be obtained from mouse long bone tissue with enzymatic digestion and fluorescence activated cell sorting. These cells can then be induced to differentiate to osteoblasts either in two-dimensional culture or following transplantation into irradiated host mice [5, 6]. Yet, unlike other perivascular stromal cell populations in the bone marrow [7, 8], the *in vivo* fate of PαS cells have not been determined during skeletal development or repair.

To determine the stromal cell populations involved in osseointegration, we used a previously developed mouse model of tibial implant surgery that mimics human joint replacement surgery (Figure 1a [9]). Following implant surgery, we performed lineage tracing experiments using tamoxifen-inducible alleles, as well as bulk and single cell transcriptome sequencing. Herein, we report significant enrichment for PαS-lineage cells at the bone-implant interface and transcriptional changes during the early stages of post-surgical healing that suggest new strategies for enhancing osseointegration.

**Figure 1:**
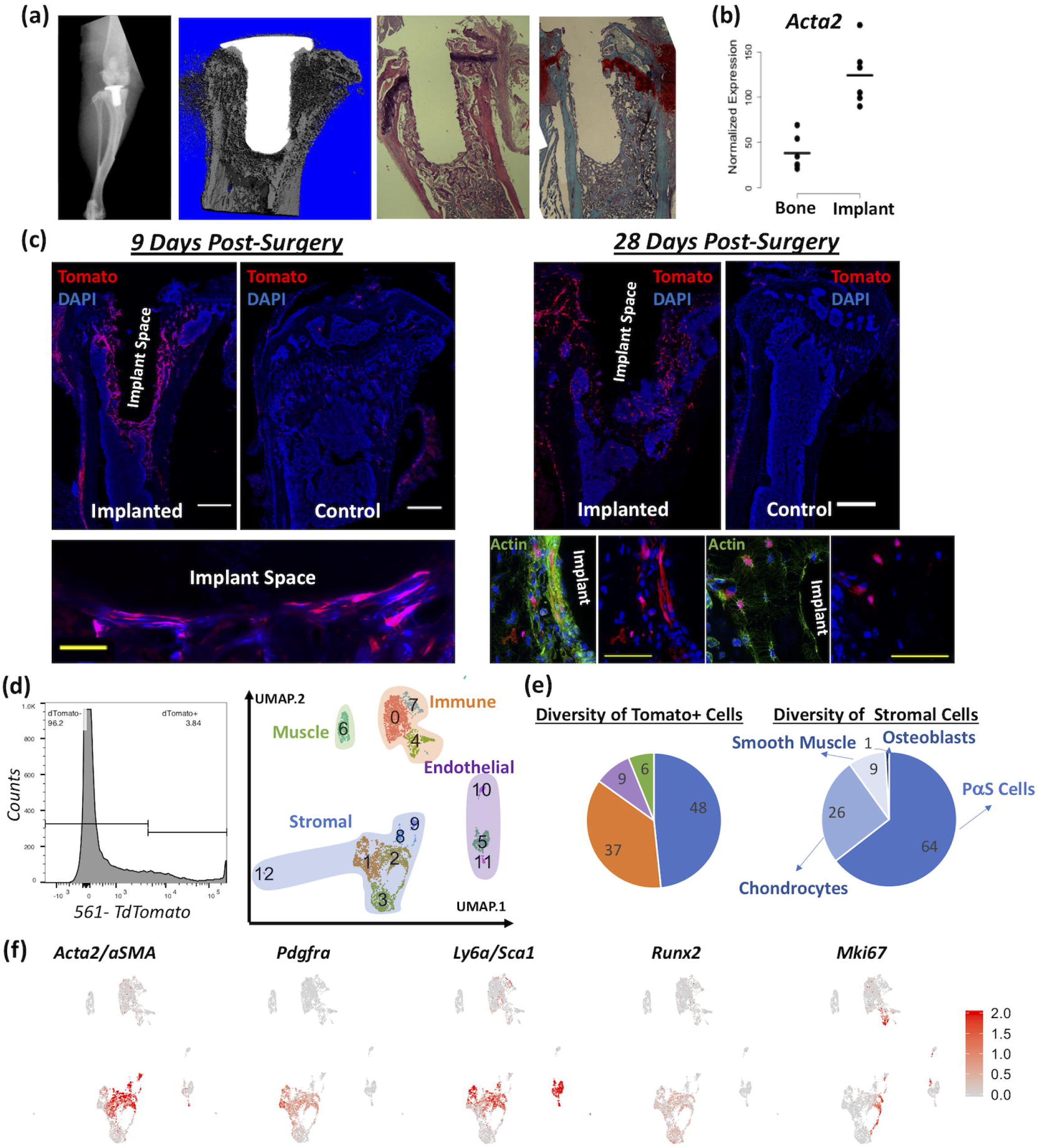
Acta2-lineage cells populate the peri-implant area following tibial implantation surgery. (a) Faxitron, microCT, hematoxylin & eosin (H&E)- and safranin O-stained images of the implanted mouse tibia are depicted (from left to right). Both microCT and H&E images indicate bone formation around the implant, while red safranin O-staining indicates lack of cartilage formation in the peri-implant area. (b) Bulk RNA-seq of peri-implant tissue from implanted tibiae indicates a significant increase in *Acta2* expression relative to cancellous bone 7 days post-surgery. (c) Implant surgery was performed on Acta2-lineage reporter mice at 20 weeks of age, with one-time tamoxifen injection immediately post-surgery. Nine days later, flat and elongated cells in the immediate vicinity of the implant are tdTom+, whereas little to no tdTomato expression was detected in the surgery-naïve control tibia (white scale bar: 500μm, yellow scale bar: 50μm). Twenty-eight days later, relatively less tdTomato expression is observed, but high magnification imaging of phalloidin-stained sections reveal tdTom+ osteocytes in the peri-implant environment (Representative images of n=3 mice/group depicted.). (d) Nine days post-surgery, tdTom+ cells from implanted tibiae were purified with flow cytometry and analyzed with single cell RNA-seq. Stromal, immune, muscle and endothelial cell populations were identified. (e) Stromal cells constituted the largest tdTom+ cell group, while the majority of them (64%) were Pdgfra+ & Sca1+ (PαS) cells. (f) Heatmaps of gene expression indicate distinct markers for each cell population. *Acta2, Pdgfra* and *Ly6a/Sca1* are co-expressed by the majority of stromal cells. Lower levels of *Runx2* expression is also detectable in the same group of cells.

## METHODS

### Mice

All experiments were approved by the Institutional Animal Care and Use Committee of Weill Cornell College of Medicine. C57/BL6J (Stock#: 000664), *Tg(Tmem100-EGFP/cre/ERT2)30Amc/J* (Stock#: 014159, hereafter referred to as *Tmem100.creERT2*) and *Gt(ROSA)26Sor*^*tm14(CAG-tdTomato)Hze*^*/J* (Stock#: 007914, hereafter referred to as *Ai14.R26.tdTomato*) mice were purchased from Jackson Laboratories (Bar Harbor, ME). *Acta2.creERT2* mice were previously described[10]. Using the aforementioned strains, we generated *Tmem100.creERT2; Ai14.R26.tdTomato* and *Acta2.creERT2; Ai14v.R26.tdTomato* mice for lineage tracing experiments. All mice were maintained in standard housing conditions, genotyped at 14 days and weaned at 21 days. To induce Cre-mediated recombination, we administered tamoxifen (Sigma cat#T5648, dissolved in corn oil) once to each mouse intraperitoneally, either at P10 (0.4mg) or at the time of implant surgery between 16-20 weeks of age (2mg). Mice were euthanized with exposure to CO_2_ at the indicated time-points.

### Tibial Implantation Surgery

Surgeries were performed as previously described[9]. Briefly, mice were anesthetized with isoflurane inhalation. The fur around the right knee joint was shaved, and skin was sterilized with betadine and chlorhexidine. Bupivacaine was injected subcutaneously for local anesthesia. An 8-mm midline incision was made over the knee. A medial para-patella incision was made to laterally dislocate the patella. The anterior cruciate ligament was excised, followed by the removal of both menisci and trimming of the tibial plateau. A hole was made by a 0.9-mm burr into the medullary canal through the tibial plateau. A 3D-printed titanium implant was press-fit into the hole. The joint was irrigated with PBS and the arthrotomy and the skin were closed with sutures. Mice received meloxicam (2mg/kg, immediately post-surgery) and buprenorphine (0.5mg/kg subcutaneously, every 12 hours for 3 days) for pain relief. Operated mice ambulate without obvious pain immediately after recovery from anesthesia. The implant articulates with femoral condyles and bears weight, similar to human joint replacements.

### Bulk RNA-seq

Tibial specimens with the implant were collected immediately after euthanasia and quickly cleaned of surrounding soft tissue. Each implant was gently removed with forceps, placed in an individual sterile plastic tube and snap-frozen in liquid nitrogen. Then, using a 1mm biopsy punch, metaphyseal cancellous bone was collected from each tibial specimen and transferred to an individual plastic tube to be frozen. Total RNA was extracted from individual peri-implant tissue and cancellous bone specimens with phenol-chloroform separation followed by on-column purification. Total RNA was then used in library preparation with the Illumina TruSeq Kit (Illumina, San Diego, CA). The libraries were sequenced on the Illumina HiSeq 4000 platform. Raw reads were mapped to the mouse genome with STAR [11] and analyzed with custom R scripts[12].

### Flow Cytometry & Single Cell RNA-seq

For evaluation of intact bone tissue, tibiae and femora from each mouse were removed at the indicated timepoints immediately after euthanasia. The epiphyses were cut with scissors, and marrow was flushed with centrifugation. For evaluation of surgically treated tibiae, the proximal half of the tibia was dissected (without disturbing the implant) and centrifuged to remove marrow. In both cases, the specimens were then collected in a sterile mortar inside a tissue-culture hood, gently crushed with a pestle, and transferred to a 15ml tube with 8ml collagenase solution (∼4,000 units in αMEM with 1% anti-mycotic; Type IV, Worthington Biochemical Corp, Lakewood, NJ). The solution containing the tissue fragments was then continuously agitated for 1 hour at 37°C. Cells were collected with centrifugation, re-suspended in αMEM (with 10% FBS and 1% anti-mycotic) and then processed on an Aria II or Influx flow cytometer (BD Biosciences, San Jose, CA) to separate tdTomato-expressing (tdTom+) cells (with gates set-up with respect to negative control cells from tamoxifen-naïve mice). The sorted tdTom+ cells were then transferred to the 10X Chromium platform to be captured in oil droplets and barcoded with cell-specific oligonucleotides. Single cell RNA-seq libraries were prepared according to the manufacturer’s instructions (10X Genomics, Pleasanton, CA) and sequenced on the Illumina HiSeq 4000 platform (Illumina, San Diego, CA). The sequencing data were processed with the Cellranger and Seurat (v3 [13]) data analysis pipelines. Trajectory analysis was performed with Monocle v2, using the standard semi-supervised algorithm[14]. Gene set enrichment analysis was performed using previously published and publicly available software [15].

### Fluorescent Histology

Following euthanasia, bone specimens were carefully removed and placed in ice-cold 4% paraformaldehyde, with gentle agitation at 4°C overnight. The specimens were washed with cold PBS and transferred to 0.5M EDTA for decalcification (for 2-4 days depending on the age of the mouse). The specimens were then incubated inside 30% sucrose until they sank. The implants were gently removed from the tibia with forceps, and the bone specimens were embedded in OCT. 20μm-thick sections were cut with a microtome (Leica Biosystems, Buffalo Grove, IL) and counter-stained with DAPI. The slides were imaged with a confocal microscope (LSM 880, Zeiss, Oberkochen, Germany) to collect images at 20 planes at 0.5μm distance from each other, using identical settings for each specimen. The fluorescent images were adjusted for brightness and contrast with Fiji/ImageJ. n≥3 specimens were evaluated per group and timepoint in each experiment.

### Phalloidin Staining

Frozen tissue sections were sequentially washed with PBS (20min), 0.3% Triton (15min), PBS (5min) at room temperature and stained with Alexa Fluor 488-conjugated phalloidin (Thermo Fisher, cat#A12379) overnight at 4°C. The next day, specimens were washed with PBS (3x, 5min each), stained with DAPI (Sigma, cat#10236276001) for 10min, re-washed with PBS (2x, 5min each), and mounted with coverslip. Specimens were stored at 4°C until imaging the following day.

### Hematoxylin-Eosin & Safranin-O Staining

Tibial specimens were fixed in 10% formalin at room temperature for 4 hours and decalcified in 0.5M EDTA for 4 days. The implants were removed with forceps and the bones were embedded in paraffin. Each tissue block was sectioned at 7μm thickness, deparaffinized with xylene, rehydrated with ethanol, and incubated in hematoxylin and eosin or safranin O solutions. The slides were further washed with ethanol and xylene, mounted with a coverslip and imaged with a brightfield microscope (Eclipse 50i, Nikon, Tokyo, Japan).

### Fluorescence Activated Cell Sorting (FACS) of PαS Cells

Cells were collected from hindlimb bone tissue as described under Flow Cytometry & Single Cell RNA-seq, resuspended in buffer solution (PBS + 2% FBS + 2mM EDTA) and blocked with recombinant Fc protein (1:100 dilution, BD Biosciences, cat#553142). Cells were then incubated with the primary antibody solution (1:100 dilution; PDGFRA-APC, Thermo Fisher, cat#17-1401-81; SCA1-FITC, Thermo Fisher, 11-5981-82; CD45-BV450, BD Biosciences, cat#560697; TER119-BV450, BD Biosciences, cat#560504; CD31-Pacific Blue, BioLegend, 102422; DAPI, Sigma, cat#10236276001) for 30 minutes on ice, washed with buffer solution and analyzed using the BD FACSCanto system (BD Biosciences, San Jose, CA). Specimens were analyzed in triplicate with at least 100,000 events per replicate.

## RESULTS

### Acta2 expression marks stromal cells around metal implants in vivo

We performed bulk RNA-seq on peri-implant tissue and control cancellous bone tissue to identify transcripts that significantly differed between them. Tibial implants were inserted into 16-week-old C57/BL6 male mice and removed 7-days later. We found significant differences in the abundances of 1173 transcripts (FDR<0.05, fold change>2), with 310 enriched in peri-implant tissue compared to cancellous bone. Within these 310 genes, we observed that *Acta2* (encoding alpha smooth muscle actin, i.e. aSMA) transcripts were enriched >3-fold (Figure 1b). Since increased *Acta2* expression has previously been reported during endochondral fracture healing in mouse long bones and *Acta2*-expressing cells can differentiate to osteoblasts and osteocytes [10, 16], we hypothesized that *Acta2*-expressing stromal cells are also involved in osseointegration. We tested this hypothesis by performing Acta2-lineage tracing. We generated *Acta2.creERT2*;*Ai14.R26.TdTomato* mice (i.e. *Acta2*-lineage reporter mice), performed tibial implant surgery at 20-weeks of age, administered a single tamoxifen dose at the end of surgery, and imaged the implanted and contralateral control tibiae for tdTomato expression 9 days later (Figure 1c). We observed abundant tdTom+ cells largely confined to the peri-implant area in the implanted tibia, and only a few tdTom+ cells on the control tibia (Figure 1c). We then repeated the experiment but evaluated tibial specimens 28-days after surgery. There remain a large number of tdTom+ cells in the implanted tibia, with many now appearing to have become osteocytes embedded in bone tissue, as indicated by phalloidin staining (Figure 1c). These results indicate that *Acta2*-expressing cells are progenitors for cells that participate in osseointegration.

To better characterize the *Acta2*-expressing cells, 9 days after surgery we recovered tdTom+ peri-implant cells from the Acta2-lineage reporter mice by sequential enzymatic digestion followed by flow cytometry, and subjected the sorted cells to single cell RNA-seq (Figure 1d). Pooling cells from n=4 implanted mouse tibiae enabled us to sequence the transcriptomes of n=4,397 cells. Clustering analysis identified 4 major groups of tdTom+ cells: Stromal cells (*Col1a1*+, 48%), leukocytes (*Ptprc/*Cd45+, 37%), skeletal muscle cells (*Acta1*+, 6%) and endothelial cells (*Pecam1*/Cd31+, 9%) (Figure 1e). Within the stromal cell clusters, the top 2 most abundant sub-populations (#1 and #2) together accounted for 64% of cells and expressed *Pdgfra* and *Sca1*, in addition to *Acta2* (Figure 1f).

*Pdgfra* and *Sca1* are expressed by a pericyte population (PαS cells) in the mouse bone marrow; these cells are capable of self-renewal and bone formation when cultured or transplanted to another mouse [6]. PαS cells are thought to be dormant during skeletal homeostasis [6], but their fate after skeletal injury has not been determined with lineage tracing. Because the PαS cells we identified in our model also express pro-osteogenic genes (such as *Runx2* and *Pth1r*) and other transcripts previously associated with fracture repair (such as *Acta2* and *Dkk3*), whereas PαS cells from uninjured bone tissue express these transcripts at very low or undetectable levels (as measured by bulk RNA-seq [17]), we decided to lineage trace PαS cells following tibial implant surgery. As *Acta2* expression is induced in PαS cells only during bone repair, the *Acta2.creERT2* transgenic mouse model cannot be used to efficiently label or modify PαS cells in uninjured bone tissue. We therefore searched for inducible Cre-expressing mouse strains that could be used to trace PαS cell activity *in vivo*, in the absence as well as presence of bone injury. Unfortunately, existing alleles associated with the principal markers of PαS cells were not suitable. *Pdgfra.creERT2* mice have poor recombination in long bone tissue [18]. Similarly, *Sca1.MerCreMer* allele have poor recombination in the long bones and vertebrae ([19], Supplementary Figure 1). We therefore screened a single cell RNA-seq dataset we previously obtained using uninjured mouse endocortical long bones [20]) for other transcripts that may mark PαS cells. We found that the expression of *Tmem100* (encoding transmembrane protein 100, previously reported to be expressed by endothelial cells[21] and neurons[22]) is largely limited to *Pdgfra*+ & *Sca1*+ cells in endocortical bone (as well as a smaller number of osteoblasts, Supplementary Figure 2). Thus, Tmem100-expression may be used to label and trace PαS cells *in vivo*.

### Postnatal Tmem100.creERT2 expression marks PαS cells, osteoblasts and endothelial cells

To determine if we can utilize *Tmem100*-transgenic mice in studying PαS cells, we obtained BAC transgenic *Tmem100*.creERT2 mouse (originally developed by Dr. Andrew McMahon) from Jackson Laboratory. We crossed male *Tmem100.creERT2* mice to female *Ai14v.R26.tdTomato* mice in order to generate *Tmem100*-lineage reporter mice. Before performing tibial implant surgery on *Tmem100*-lineage reporter mice, we characterized the specific skeletal cell populations in which *Tmem100*-transgene recombines upon tamoxifen administration. Following a single tamoxifen injection at P10, we evaluated tibial and spinal specimens at 3-, 8-, 30-, 120- and 180-days post-injection. We found extensive tdTom+ cells along cortical bone surfaces (endosteum in particular) and the primary spongiosum 3-days later (Figure 2a and Supplementary Figure 3). We also gave Tmem100-lineage reporter mice a single dose of tamoxifen at P60 (data not shown) or P120 (Figure 3b); again, we observed tdTom+ cells along bone-lining surfaces, indicating that the *Tmem100*-transgene is expressed by bone-lining cells in neonates and in adults. Interestingly, we were unable to detect GFP-expression with histology or flow cytometry.

**Figure 2:**
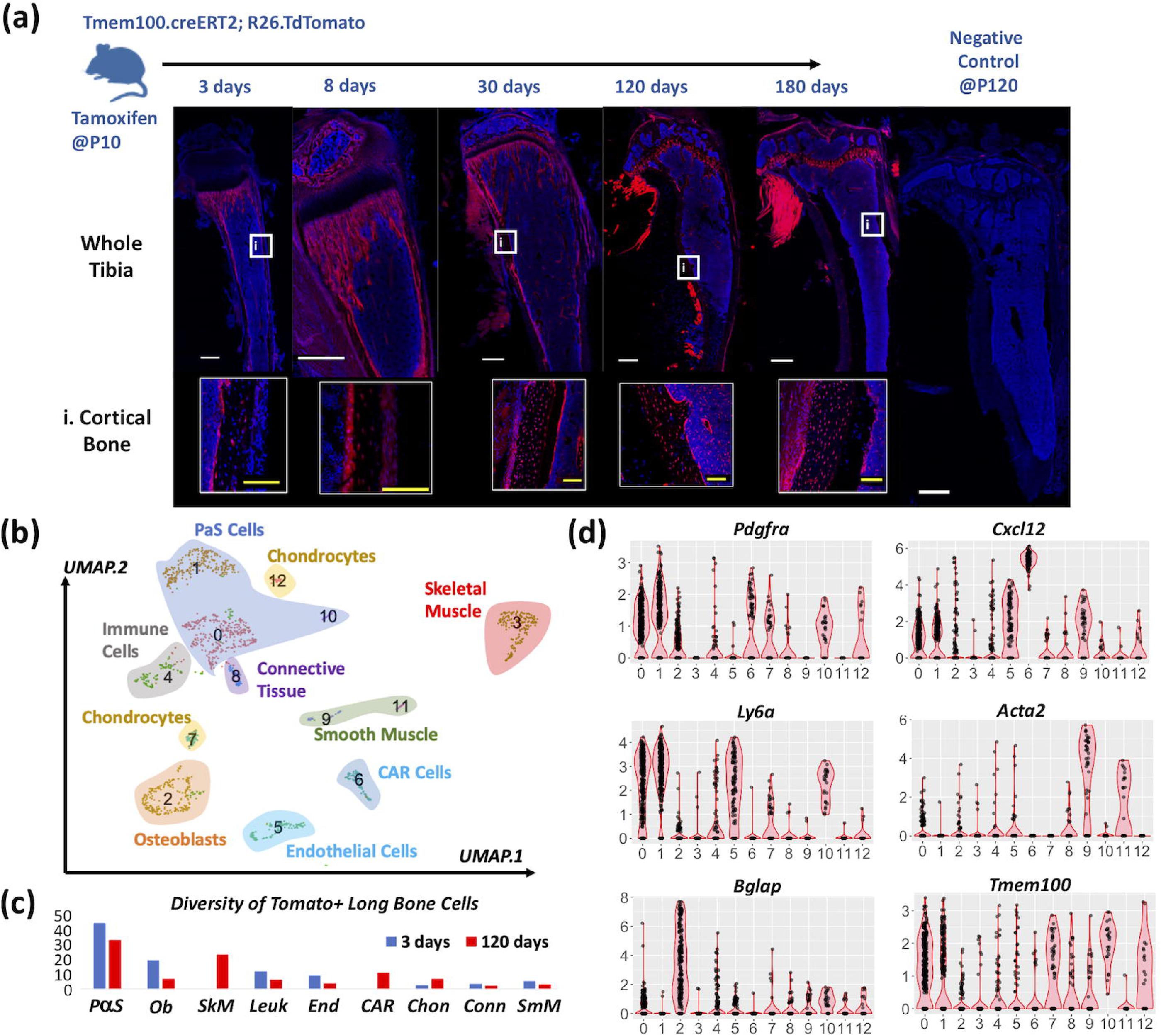
Tmem100.creERT2 transgene labels PαS cells in the developing skeleton. (a) Tmem100-lineage reporter mice were injected with tamoxifen at P10 and evaluated at 3-, 8-, 30-, 120- and 180-day time points post-injection. The proportion of tdTom+ osteocytes in the diaphyseal cortex increased over time (Representative images of n≥3 mice/group depicted. White scale bar: 500μm, yellow scale bar: 100μm). (b) Single cell RNA-seq of tdTom+ cells at 3- and 120-days indicates that approximately 40% of labeled cells are PαS cells (represented by clusters #0 and #1) at both time-points. (c) The transcriptional diversity of Tmem100-lineage cells were similar at 3- and 120-days post-injection, with some differences. >20% of the cells detected at 120-days were contaminating skeletal muscle cells; these cells were not captured at the 3-day timepoint. Cxcl12-abundant reticular (CAR) cells are also observed at 120-days but not at 3-days post-injection. (d) Violin plots depict the expression of cluster-specific markers. Clusters #0, #1 and #10 co-express Pdgfra and Ly6a/Sca1, whereas clusters #9 and #11 express Acta2, likely representing smooth muscle cells.

**Figure 3:**
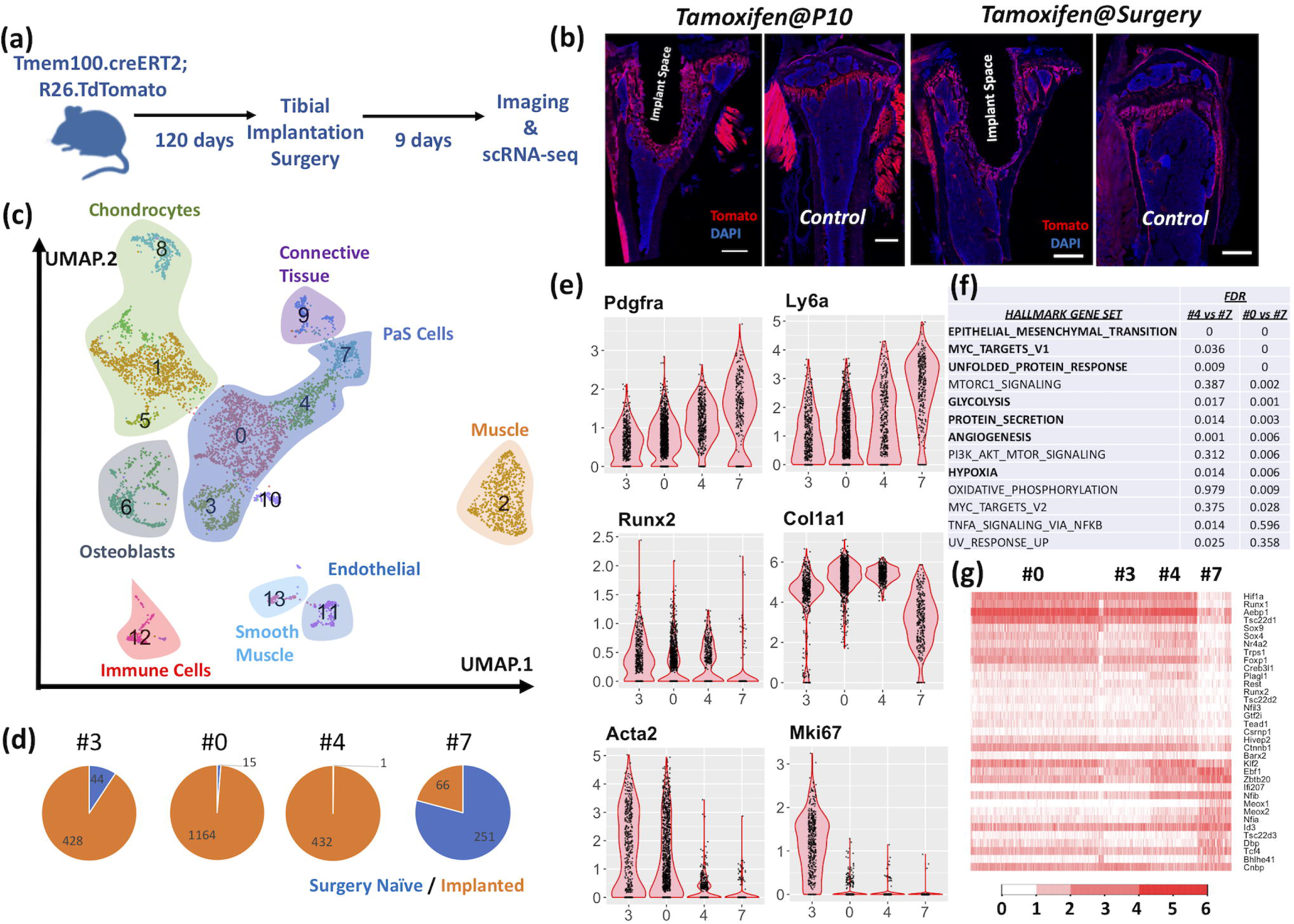
Implantation surgery on Tmem100-lineage reporter mice results in labeling of peri-implant cells similar to Acta2-lineage reporter mice. (a) Tmem100-lineage reporter mice were subjected to tibial implantation surgery at 4 months. (b) Mice were injected with tamoxifen either at P10 (left) or at the time of surgery. Regardless of the timing of tamoxifen treatment, peri-implant cells were labeled with tomato expression (Representative images of n≥3 mice/group depicted. Scale bar: 500μm). (c) tdTom+ cells were purified from the implanted or surgery-naïve tibiae of mice treated with tamoxifen at surgery and evaluated with single cell RNA-seq. Clusters #0, #3, #4 and #7 were found to express markers of PαS cells. (d) Pie charts indicate little overlap between the PαS cells originating from implanted and surgery-naïve tibiae of Tmem100-lineage reporter mice. (e) Violin plots show that PαS cells collected from implanted tibiae express Acta2, Runx2 and high levels of Col1a1, whereas the cells from surgery-naïve tibiae do not. (f) Gene set enrichment analysis (GSEA) identified significant enrichment for multiple groups of genes, when clusters #0 and #4 are separately compared to #7. Gene sets that are significantly enriched in both comparisons (FDR<0.05) are highlighted in bold. (g) Heatmap of transcription factors with significant changes in expression across clusters associated with PαS cells.

To determine whether PαS cells were tdTom+, we separated PαS cells from the long bones of P13 mice (that received tamoxifen at P10) by flow cytometry (Pdgfra^+^, Sca1^+^, Cd45^-^, Cd31^-^, Ter119^-^) and found that >80% were tdTom+ (Supplementary Figure 4); thus, *Tmem100*-transgene is expressed by most PαS cells in long bones. To determine which cell populations in addition to PαS cells express the *Tmem100*-transgene, we performed single cell RNA-seq on tdTom+ cells recovered from long bones and vertebrae of *Tmem100*-lineage traced mice, at 3 days and 120 days after tamoxifen injection at P10 (Figure 2b). Forty percent of all tdTom+ cells were PαS cells. Other tdTom+ cells included endothelial cells (*Pecam1*/*Cd31*+, *Emcn*+, 7%) osteoblasts (*Bglap*+, 14%), leukocytes, and skeletal muscle cells (Figure 2c). The relative proportion of tdTom+ cells that were PαS cells was comparable between long bone and spine specimens and did not change substantially between the 3-day and 120-day post-tamoxifen specimens. We detected high *Acta2*-expression in 2 small cell clusters (#9 and #11, Figure 2c-d); however, these cells did not express markers of PαS cells. Instead, their transcriptomes were consistent with those of smooth muscle cells (e.g., *Myh11*+, *Notch3*+, *Tagln*+)[23, 24] (Figure 2d).

Notably, we detected cells matching the transcriptional profile of *Cxcl12*-abundant reticular cells (i.e. CAR cells) 120-days post-tamoxifen injection, but not at 3-days (Figure 2c). Consistent with this observation, starting at 30-days post-tamoxifen injection, we detected tdTom+ cells with reticular morphology around bone marrow sinusoids (Supplementary Figure 3). Taken together, our data indicate Tmem100.creERT2 is expressed in multiple cell types with PαS cells being the most common cell type in bone. Therefore, Tmem100.creERT2 mice can be used to lineage-trace PαS cells *in vivo*.

### Post-surgical PαS cell activation is associated with significant transcriptional changes

To identify the changes in PαS cell transcriptome during implant osseointegration, we performed tibial implant surgery on Tmem100-lineage reporter mice, and compared surgically implanted tibial specimens to uninjured (control) tibiae. We first determined whether the timing of tamoxifen injection influences the spatial distribution of tdTom+ cells in the peri-implant area. We pulsed one group of mice with tamoxifen at P10 and performed surgery on them 4 months later, whereas we kept another group of mice tamoxifen-naïve and pulsed them with tamoxifen at the end of surgery at 4 months of age (Figure 3a-b). Nine-days post-surgery, we observed that peri-implant cells in both groups of mice were tdTom+, indicating that cell-type-specific Tmem100-transgene expression patterns are similar at P10 and P120.

We next collected tdTom+ cells from the surgically treated tibiae and intact tibiae of mice (separately pooled from n=4 mice) treated with tamoxifen after surgery, and performed single cell RNA-seq (Figure 3c). As expected, PαS cells were abundantly present in the collective single cell RNA-seq data, however cells from implanted and surgery-naïve control tibiae separated into distinct clusters, indicating significant differences between their transcriptomes. The tdTom+ cells retrieved from implanted tibiae contained PαS cells (clusters #3, #4 and #0) that expressed markers of proliferation (e.g. *Mki67* in cluster #3) and osteogenic differentiation (e.g. *Runx2*) whereas those from the control tibiae (cluster #7) did not (Figure 3d-e). To determine the transcriptome-wide differences between PαS cells from implanted and control tibiae, we compared the cells in clusters #4 and #0 with those in cluster #7, using Seurat’s FindMarkers function [25]. We found that 647 genes were differentially expressed, with 498 upregulated and 149 downregulated genes in both clusters #0 and #4, compared to #7 (adjusted p<0.05, Supplementary Table 1). The upregulated genes included *Col1a1, Col1a2, Runx2, Pth1r, Ibsp* and *Spp1* (Figure 3e), altogether suggesting that PαS cells differentiate towards an osteogenic fate following tibial implant surgery. To further confirm this, we performed a trajectory analysis of all PαS cells (clusters #0, #3, #4 and #7) and osteoblasts (cluster #6). Monocle positioned quiescent PαS cells from control tibiae and osteoblasts from both groups of tibiae at the opposite ends of the pseudotime spectrum (Supplementary Figure 5). However, PαS cells from the implanted tibia were positioned along the trajectory that connects the aforementioned cell types (Supplementary Figure 5), further indicating that implant surgery induces differentiation of dormant PαS Cells.

To determine if the differences between quiescent and surgery-activated PαS cells might be indicative of specific molecular processes, we performed a gene set enrichment analysis (GSEA). We found significant changes in multiple sets of genes related to cellular differentiation and stress (e.g. epithelial-mesenchymal transition, unfolded protein response), metabolism (e.g. glycolysis), immunity (e.g. Tnfa-signaling) and development (e.g. angiogenesis) (Figure 3f) as a result of implant surgery. When we evaluated known transcription factors that might be involved in activation of quiescent PαS cells, we observed increased expression of 22 transcripts (including multiple factors previously associated with skeletogenesis, such as *Creb3l1[26], Ctnnb1[27], Nr4a2[28], Runx1[29], Sox4[30]* and *Sox9[31]*, in addition to *Runx2[32]*), as well as decreased expression of 13 transcripts (including *Ebf1[33], Id3[34]* and *Tsc22d3[35]*) in clusters #0 and #4 (Figure 3g). These results indicate that tibial implant surgery induces a highly diverse set of transcriptional changes in PαS cells in addition to induction of differentiation.

## DISCUSSION

Peri-articular titanium implants are commonly utilized in orthopaedic procedures, such as total knee and hip joint replacements. Robust osseointegration is critical to the longevity of these implants. However, the molecular and cellular mechanisms of osseointegration are unclear. Here, we show that the insertion of a titanium implant into the mouse tibia activates PαS cells in the peri-implant region. While PαS cells are quiescent in uninjured bone, our data indicate that these cells proliferate and undergo osteogenic differentiation in newly formed peri-implant bone tissue (summarized in our model in Supplementary Figure 6). We also find that the activation of PαS cells is associated with a transcriptional signature that includes increased expression of osteo-anabolic, metabolic, stress-induced and immunity-related genes (Figure 3f). One interesting increase involves *Pth1r* (encoding parathyroid hormone receptor), whose expression was not observed in dormant PαS cells. Whether implant surgery-activated PαS cells are more responsive to parathyroid hormone (PTH) therapy requires formal testing. However, consistent with this hypothesis, PTH treatment has previously been shown to significantly increase the biomechanical strength of bone-implant interface in our model[9].

Additional molecular mechanisms that follow PαS cell-activation still need to be delineated. Intriguingly, one transcription factor that is downregulated in implant-activated PαS cells is Early B cell factor, *Ebf1*. EBF1 is a negative regulator of bone mass [33, 36, 37]. However, the cell types responsible for this phenotype has not been identifed, as conditional deletion of *Ebf1* with *Lepr-Cre* or *Runx2-cre* did not alter bone properties[36, 37], whereas global or *Prrx1-Cre*-mediated deletion of *Ebf1* increased trabecular bone mass [33, 38]. It will be important to determine if conditional inactivation of *Ebf1* in PαS cells leads to improved bone properties.

Quiescent PαS cells do not express the *Acta2* transcript at levels detectable by single cell RNA-seq (Figure 3d). Our data indicate that implant surgery leads to *Acta2.creERT2* expression in PαS cells. Our findings are consistent with reports that performed *Acta2*-lineage-tracing in other models of bone repair, such as endochondral fracture healing [10, 16] and anterior cruciate ligament reconstruction [39]. Thus, *Acta2* expression is highly dynamic in healing bone tissue and is induced in PαS cells immediately after injury.

Because *Acta2.creERT2* expression occurs after PαS have been activated, our discovery that *Tmem100.CreERT2* expression precedes or is contemporaneous with PαS formation provides a new tool for manipulating PαS cells *in vivo. Tmem100.CreERT2* is expressed in PαS cells and osteoblasts; therefore, this transgene may not be appropriate for developmental osteogenic fate-mapping studies. Nevertheless, *Tmem100.CreERT2* is expressed in a large fraction of PαS cells and could be used to temporally modify these cells. Moreover, the single cell RNA-seq data we generated suggests there are other markers that can be used to define PαS cells and distinguish quiescent cells from implant-activated ones (Supplementary Table 1). These additional markers are particularly important since *Ly6a*, which encodes Sca1, does not have a known human ortholog. Therefore, while mice have PαS cells, their human counterpart remains to be identified. In this regard, *Cd248* (which encodes the transmembrane protein endosialin) is one candidate marker that warrants follow-up. Endosialin expression was restricted to PαS cells in our long bone single cell RNAseq data. In other tissues endosialin expression has been observed in pericytes found in fat, aorta and synovium, and can be utilized for prospective isolation of stromal cells with flow cytometry [40]. Humans have a CD248 ortholog.

One intriguing outcome of our experiments is the identification of tdTom+ CAR cells in the bone marrow not immediately after tamoxifen injection at P10, but starting at 30-days and later time-points (Figure 2c and Supplementary Figure 3). These data suggest the possibility that PαS cells may give rise to CAR cells, as arteriole-type large blood vessels (surrounded by pericytes including PαS cells) sprout sinusoids (surrounded by CAR cells) during the development of marrow vasculature[41]. Yet, similar observations have been reported with the *Sp7.CreERT2* mouse model[42], wherein early postnatal labeling of *Sp7.Cre*-expressing cells results in the labeling of CAR cells later on. The degree of overlap between the cells targeted by *Tmem100.CreERT2* and *Sp7.CreERT2* mouse models remains unclear. However, Cre-recombination in CAR cells has been reported in other congenitally activated mouse models, such as *Bglap.Cre* and *Dmp1.Cre* [43]. Therefore, it is also possible that a small number of tdTom+ CAR cells at P10 in Tmem100-lineage reporter mice are capable of proliferation (as suggested by Seike et al.,[36]), in order to give rise to a larger number of CAR cells later on. Development of novel mouse models with PαS cell-specific recombination profiles will likely provide the definitive answer.

In summary, we have discovered that quiescent PαS cells are induced to proliferate and differentiate during metal implant osseointegration in mice. Importantly, our data demonstrate the utility of scRNA-seq in understanding the plasticity of discrete cell populations (and identifying pertinent markers to guide orthogonal experiments) during bone healing; these studies would not be possible with tissue-level RNA-seq alone. Understanding how PαS cells promote osseointegration whether they are necessary and sufficient for this process should lead to new strategies for enhancing osseointegration in humans, thereby reducing the incidence of implant failure following joint replacement surgery.

## AUTHOR CONTRIBUTIONS

XY, LBI, MPGB and UMA conceived the experiments. AV, ESS, DGA, XY, MR, BS, YN and UMA performed data acquisition and analysis. ESS, DGA, XY, IK, LBI, MPGB and UMA contributed to data interpretation. All authors reviewed and approved the final version of the manuscript.

## ACKNOWLEDGMENTS

This study was supported by Hospital for Special Surgery institutional funds, and additional funds provided by the Tow Foundation for the David Z. Rosenzweig Genomics Center at HSS. The authors thank Drs. Matthew Greenblatt, Noriaki Ono, Henry Kronenberg and Matthew Warman for thoughtful comments.

## SUPPLEMENTARY DATA

Supplementary Figure 1: We tested the recombination efficiency of the Sca1.MerCreMer knock-in mouse model in the tibia and lumbar spine. (a) Sca1-lineage reporter mice were injected with tamoxifen at P11, and sacrificed 2 days later. Flow cytometry did not identify a meaningful number of tdTom+ cells among cells collected from tibial and femoral bone specimens (data not shown). Histologic analysis depicted tdTomato-labeling of little to no cells in the tibia (b) and spine (c).

Supplementary Figure 2: Single cell RNA-seq data derived from uninjured endocortical mouse cells depict cell-type specific expression of marker genes (Ayturk et al., doi.org/10.1101/849224). *Pdgfra* and *Ly6a/Sca1* expression is specific to cluster #6, whereas *Acta2* expression in intact mouse bone is specific to a cell population (cluster #19) that also expresses *Notch3* and *Mustn1*, but not *Pdgfra* or *Ly6a. Tmem100* expression is largely limited to cluster #6; these cells also co-express PαS cell markers *Pdgfra* and *Ly6a*/Sca1 (aSMA: alpha-smooth muscle actin; CAR: Cxcl12-abundant reticular cell).

Supplementary Figure 3: *Tmem100*.creERT2 transgene expression comprehensively marks a multitude of skeletal tissues in mice. (a) Cells lining the anterior cruciate ligament and meniscus are tdTom+ 3-days after tamoxifen injection at P10. (b) Morphologically distinct tdTom+ marrow cells appear at 30-days primarily around the sinusoid-type blood vessels in tibia, suggesting that these cells might be Cxcl12-abundant reticular (CAR) stromal cells. (c) Time-dependent tdTomato-expression patterns observed in the tibia are consistent with those in the lumbar vertebrae. Trabecular and cortical bone surfaces are comprehensively labeled at 3-days, whereas tdTomato-expression is rather limited to osteocytes and growth plate chondrocytes at 120- and 180-days. Nucleus pulposus is consistently labeled with tdTomato-expression at all time points.

Supplementary Figure 4: Representative FACS plots indicate that the majority of PαS cells are tdTom+ in Tmem100-lineage reporter mice. Live, single cells are gated based on forward and side scatter. Lineage-(CD45-, CD31-, TER119-) cells are evaluated based on PDGFRA-APC and SCA1-FITC fluorescence. tdTom+ portion of PDGFRA+ & SCA1+ cells are determined based on PE-fluorescence.

Supplementary Figure 5: Trajectory analysis with Monocle positions active PαS cells between dormant PαS cells and osteoblasts. (a) Clusters representing PαS cells (#0, 3, 4 and 7) and osteoblasts (#6), as identified by Seurat were subjected to trajectory analysis with Monocle. (b) Cells representing dormant PαS cells from uninjured control tibiae are positioned at the beginning of pseudotime, whereas osteoblasts from both control and implanted tibiae are positioned at the end of pseudotime. PαS cells from implanted tibiae that are presumably differentiating are scattered across the pseudotime trajectory, between dormant PαS cells and osteoblasts. (c) Representative transcripts that mark distinct stages of pseudotime are indicated in violin (left) scatter (center) and trajectory (right) plots in each panel. *Clec3b* expression marks the dormant PαS cells positioned at the beginning of pseudotime, whereas *Acta2* and *Mki67* expression marks active PαS cells positioned along the transitionary path and *Bglap* expression marks osteoblasts at the end of pseudotime.

Supplementary Figure 6: Model of PαS cell activation after tibial implant surgery. PαS cells begin to proliferate and differentiate following implant surgery. They exhibit a fibroblast-like morphology and express *Acta2, Runx2* and *Pth1r* at 9-days, whereas a subset of them go on to become osteocytes in newly formed peri-implant bone at 28-days.

Supplementary Table 1: Differential gene expression analysis of active and quiescent PαS cells reveals significant differences in 647 genes (adjusted p<0.05).

## Notes

### Competing Interest Statement

The authors have declared no competing interest.

