## Supplementary figures and images for "Bone Marrow Stromal Cells in a Mouse Model of Metal Implant Osseointegration"

### Supplementary Figure 1

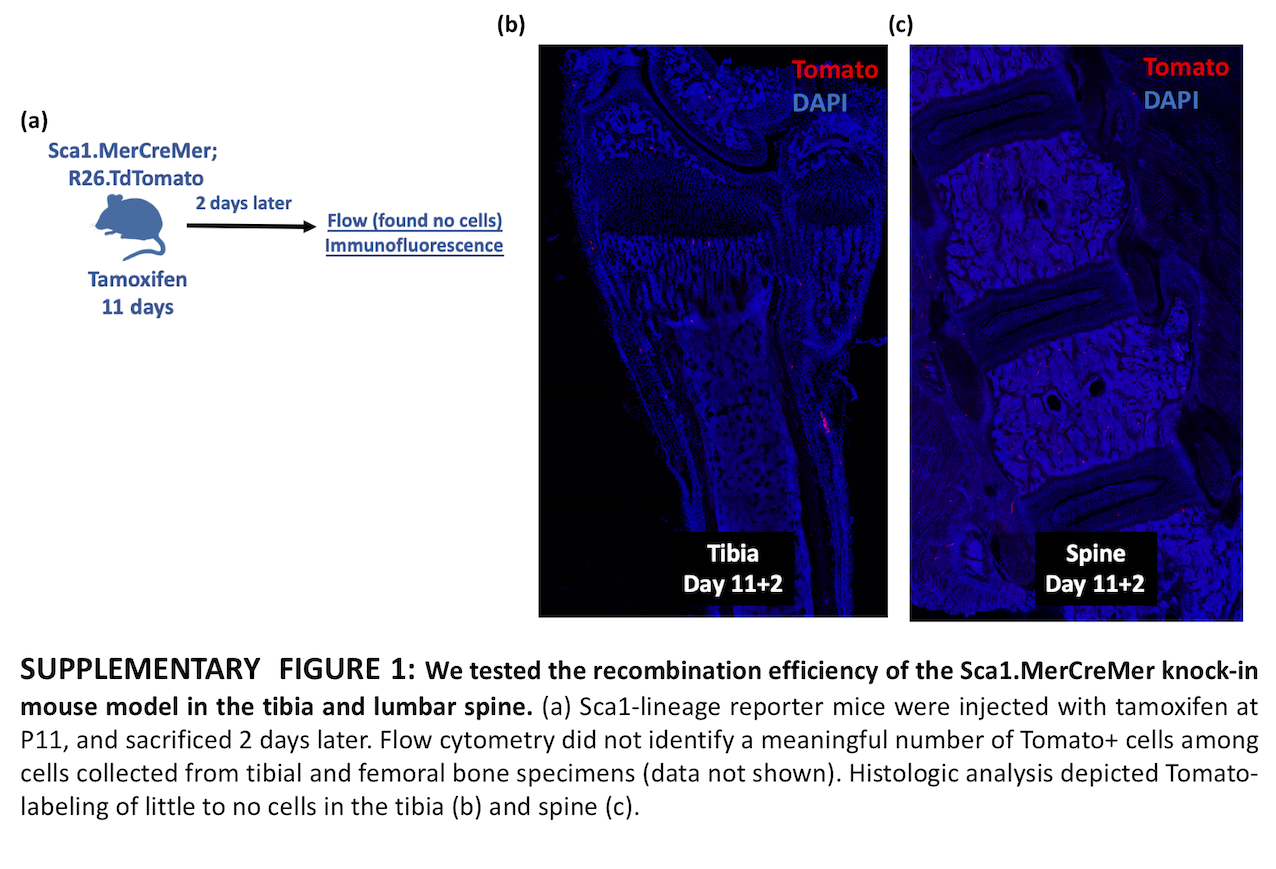

### Supplementary Figure 2

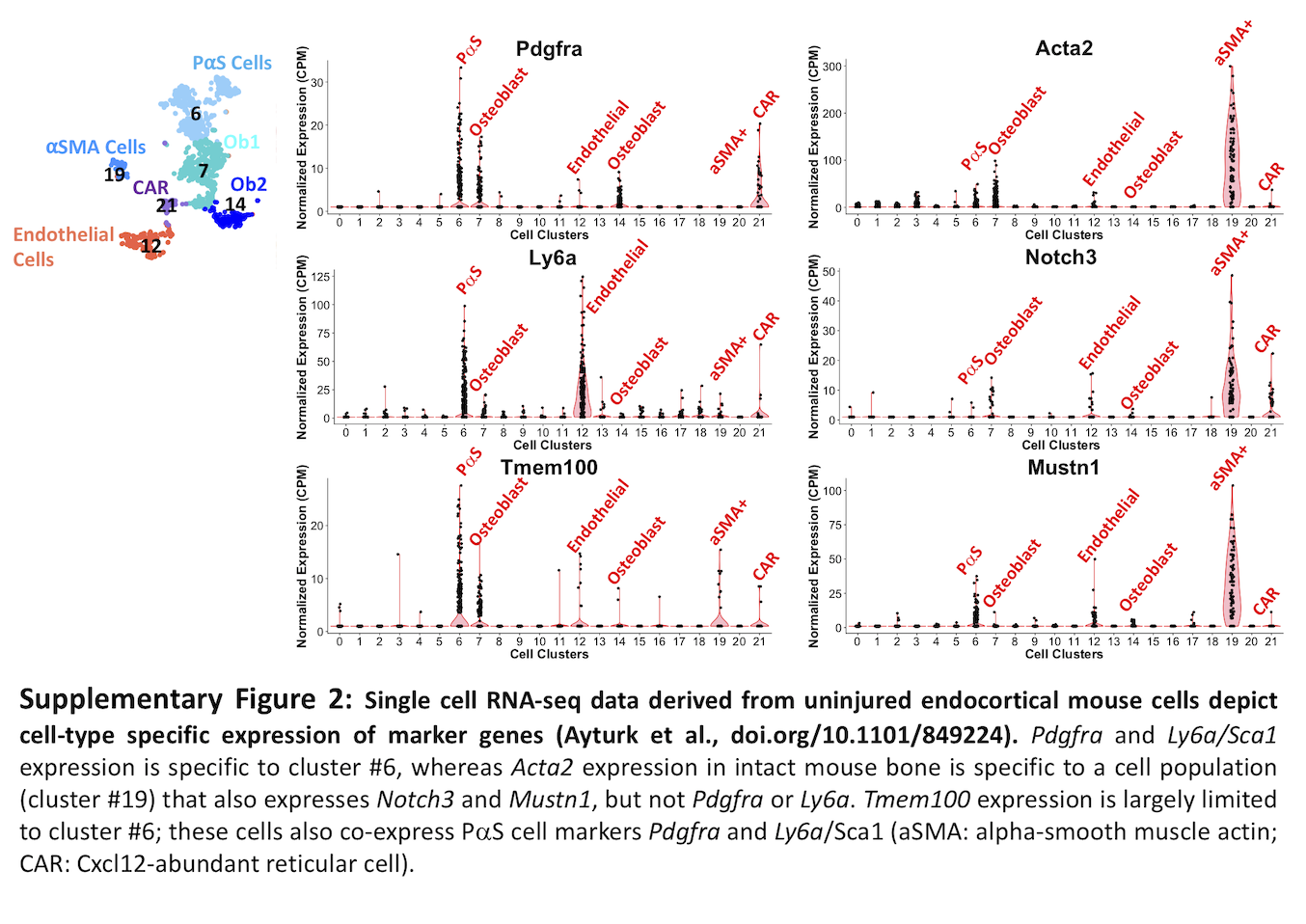

### Supplementary Figure 3

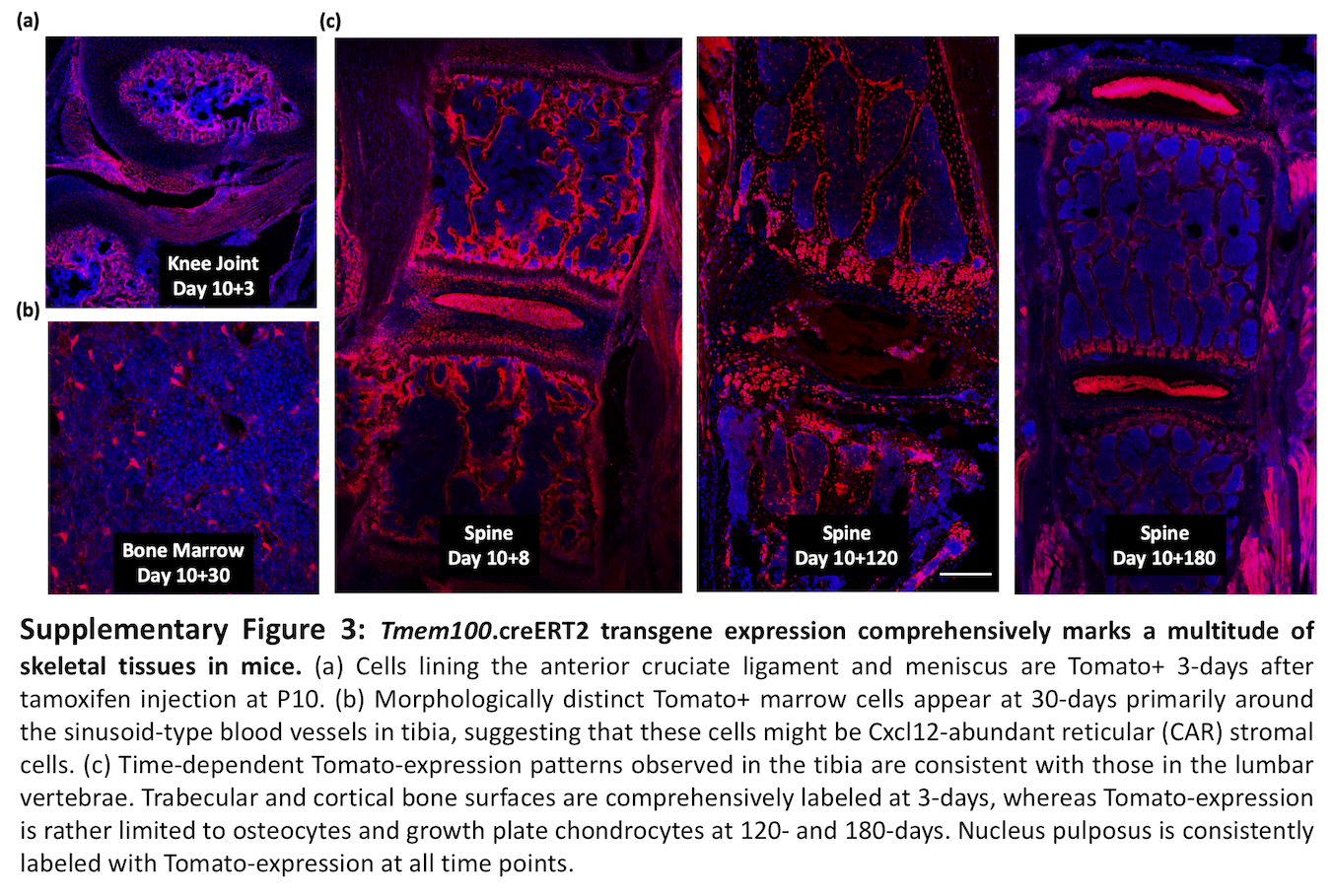

### Supplementary Figure 4

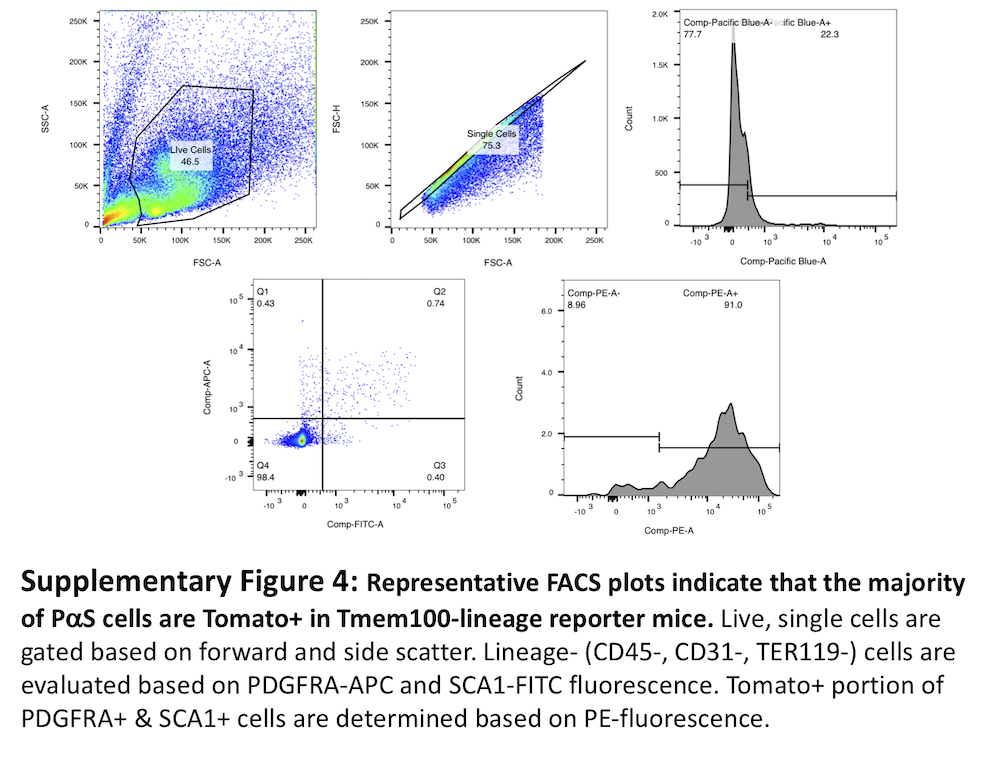

### Supplementary Figure 5

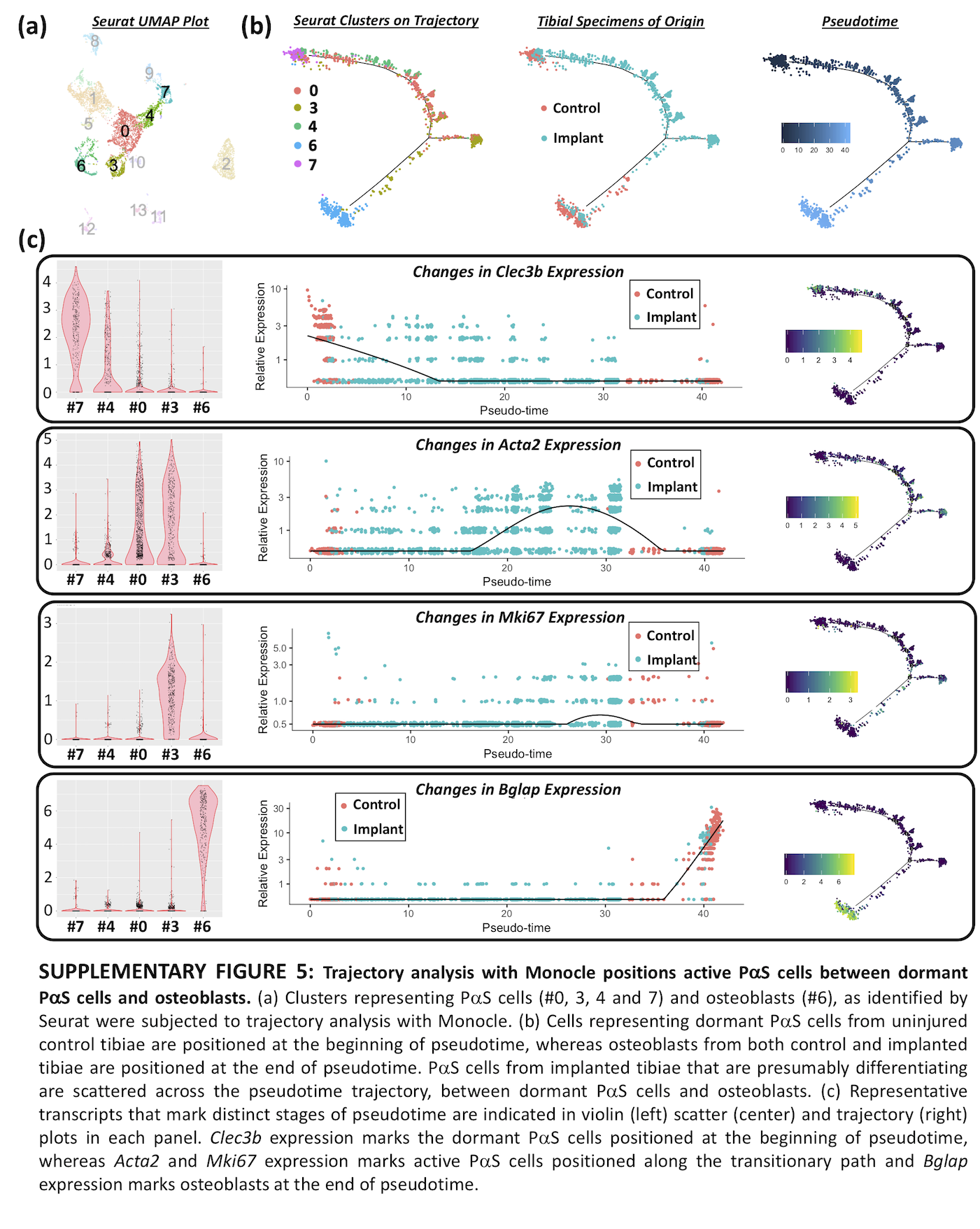

### Supplementary Figure 6

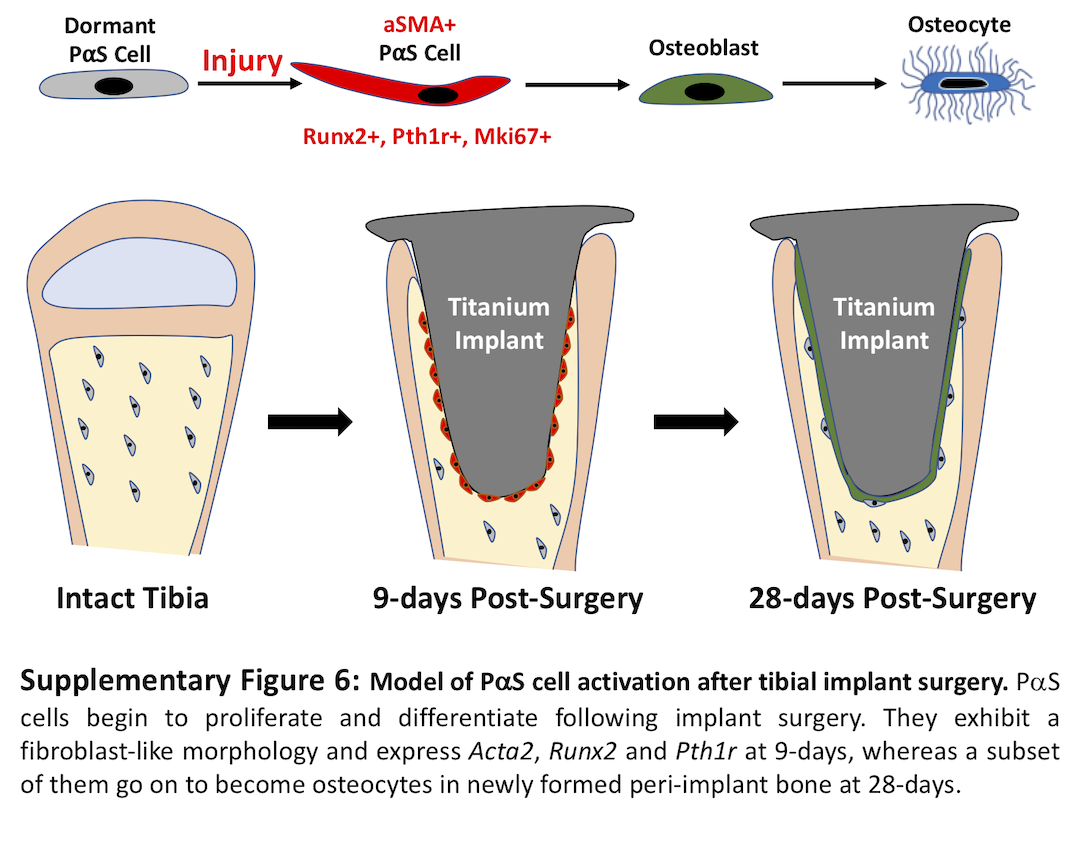
